# Intra-host variation in the spike S1/S2 region of a feline coronavirus type-1 in a cat with persistent infection

**DOI:** 10.1101/2023.07.31.551356

**Authors:** Ximena A. Olarte-Castillo, Beth. N. Licitra, Nicole M. André, Maria A. Sierra, Christopher E. Mason, Laura B. Goodman, Gary R. Whittaker

**Affiliations:** Departments of Microbiology & Immunology, Cornell University College of Veterinary Medicine, Ithaca NY, USA; Public & Ecosystem Health, Cornell University College of Veterinary Medicine, Ithaca NY, USA; Clinical Sciences, Cornell University College of Veterinary Medicine, Ithaca NY, USA; James A. Baker Institute for Animal Health, Cornell University College of Veterinary Medicine, Ithaca NY, USA; Feline Health Center, Cornell University College of Veterinary Medicine, Ithaca NY, USA; Department of Physiology & Biophysics, Weill Cornell Medicine, New York NY, USA

## Abstract

Feline coronavirus type 1 (FCoV-1) is widely known for causing feline infectious peritonitis (FIP), a systemic infection that is often fatal, with the virus known as the FIPV biotype. However, subclinical disease also occurs, in which cats may not show signs and intermittently shed the virus, including in feces, possibly for long periods of time. This virus is known as the FECV biotype. Progression of FECV to FIPV has been linked to several genomic changes, however a specific region of the viral spike protein at the interface of the spike S1 and S2 domains has been especially implicated. In this study, we followed a cat (#576) for six years from 2017, at which time FCoV-1 was detected in feces and conjunctival swabs, until 2022, when the animal was euthanized based on a diagnosis of alimentary small cell lymphoma. Over this time period, the cat was clinically diagnosed with inflammatory bowel disease and chronic rhinitis, and cardiac problems were also suspected. Using hybridization capture targeting the spike (S) gene of FCoV followed by next-generation sequencing, we screened 27 clinical samples. We detected FCoV-1 in 4 samples taken in 2017 (intestine and nasal tissue, feces, and conjunctiva), and 3 samples taken in 2022 (feces, and intestinal and heart tissue), but not in fecal samples taken in 2019 and 2020. Next, we focused on the S1/S2 region within S, which contains the furin cleavage site (FCS), a key regulator of viral transmission and pathogenesis. We show that the FCoV-1 variants obtained from feces in 2017 and 2022 were identical, while the ones from conjunctiva (2017), heart (2022), and intestine (2017 and 2022) were distinct. Sequence comparison of all the variants obtained showed that most of the non-synonymous changes in the S1/S2 region occur within the FCS. In the heart, we found two variants that differed by a single nucleotide, resulting in distinct FCS motifs that differ in one amino acid. It is predicted that one of these FCS motifs will down-regulate spike cleavability. The variant from the conjunctiva (2017) had a 6-nucleotide in-frame insertion that resulted in a longer and more exposed S1/S2 loop, which is predicted to be more accessible to the furin protease. Our studies indicate that FCoV-1 can independently persist in the gastrointestinal tract and heart of a cat over a long period of time without evidence of typical FIP signs, with intermittent viral shedding from the gastrointestinal and respiratory tracts.

## Introduction

Feline coronavirus type 1 (FCoV-1) is a common and highly transmissible coronavirus (CoV) of both domestic and non-domestic cats. FCoV-1 is a distinct virus within the Alphacoronaviruses (Jaimes et al., 2020; Whittaker et al., 2018); it is not readily isolatable in cell culture and so remains poorly studied from a virological perspective. FCoV-1 represent the majority of FCoV infections in the feline population, the prevalence of which has been estimated at 75-95% in multi-cat households (Pedersen, 1995), possibly dropping to 25% in single-cat households and approaching 100% in shelter/breeder situations. With an estimated 58 million owned cats in the US alone (according to the American Veterinary Medical Association), FCoV represents a widespread endemic coronavirus that remains largely unexplored from a molecular evolution perspective. According to recent guidelines from the American Association of Feline Practice (Thayer et al., 2022), FCoV infection has three principal clinical outcomes: animals clear what is assumed to be an acute primary infection, with no viral shedding (about 5% of cats); animals intermittently shed low levels of virus from their GI tract (70-80% of cats); animals develop long-term persistent shedding with high viral load for a prolonged period (10-15% of cats). Based on this, the virus likely persistently infects cats over long periods of time. In a subset of infected cats (about 5-12%), feline infectious peritonitis (FIP) follows the primary/persistent infection.

FIP typically presents in either an effusive (wet) or non-effusive (dry), form; typically associated with peritoneal/pleural effusion and neurological signs respectively. FIPV was originally thought to arise based on an internal mutation within the genome of the endemic feline enteric coronavirus (FECV) (Vennema et al., 1998), which allowed the virus to gain its tropism to allow productive infections of monocytic cells such as macrophages, and so cause its varied disease signs. However, the progression of FCoV (FECV), to its FIPV biotype has now been linked to several genomic changes, including in the viral 7b, 3c and spike (S) genes (Barker & Tasker, 2020; Pedersen, 2014a, 2014b). While multiple genomic changes likely account for ultimate conversion to FIP, a specific region of spike—the structural loop spanning the interface of the spike S1 and S2 domains—is linked to the FIPV phenotype. Our prior studies have shown that amino acid sequence changes in this region are highly correlated with conversion to FIP. In circulating FCoV-1, this S1/S2 domain contains a consensus motif for cleavage-activation by the cellular protease furin (*i.e*., a furin cleavage site, or FCS) (Henrich et al., 2003).

Our initial molecular analysis of S1/S2 identified a consensus sequence (S/T/Q)-R-R-(S/A)-R-R-S in 30 fecal samples from apparently healthy cats (*i.e*., FECV) and a disruption of this motif in 22 tissue samples from cats clinically confirmed to have FIP based on immunohistochemical (IHC) analysis (Licitra et al., 2013). In this initial study, the disruption of the consensus cleavage motif was present in 100% of FIP cats— although not in all tissues. Subsequent case studies of individual cats and follow up of a small FIP outbreak in an animal shelter also confirmed this 100% correlation (Andre et al., 2019; Andre et al., 2020; Healey et al., 2022). Independent validation of S1/S2 mutations as drivers of FIP has been limited, in part due to technical difficulties reported by others in sequencing this region of spike (Barker et al., 2017), although recent epidemiology studies from China have recently provided some support for this hypothesis (Ouyang et al., 2022). Notably, our recent genomic analysis has provided additional support for the “FCS” disruption hypothesis, which identified the FCS within the FCoV-1 S1/S2 loop (along with other residues) as site of natural selection between pathogenic and non-pathogenic FCoVs (Zehr et al., 2023). This genomic analysis also identified “M1058L” as a site of natural selection. “M1058L” has long been attributed to systemic spread of FCoV (but not with FIP *per se*) (Barker et al., 2017; Meli et al., 2022)

Alongside natural infections of felines, experimental infection of cats with FCoV-1 has also shown several patterns of virus shedding and intra-host variation over time. For example, infection of immunocompromised cats with an “FECV” isolate RM resulted in 2 out of 18 cats (11%) developing FIP. Sequencing of the 7b gene of the viruses isolated from these two cats revealed these new variants, named UCD-9 and UCD-10, were highly similar (89.32%) and grouped with the original RM virus; however detailed genomic analysis of these viruses has not been performed. Subsequent experimental inoculation of laboratory-housed cats with these new variants resulted in the development of FIP in 50% of the inoculated cats (Poland et al., 1996). Experimental infection of cats has also shown that even if a cat does not develop FIP over time, the S gene is a hot spot for mutation (Herrewegh et al., 1997) (Desmarets et al., 2016). In naturally infected cats, persistent infection with FCoV-1 has been shown, with viral shedding in feces detected up to seven months after the first detection in a closed-cat breeding facility (Herrewegh et al., 1997) and up to 6 years in two household cats in the UK (Addie et al., 2003). In addition to these studies, the consistent finding of FCoV-1 in the colon of cats after both experimental (Kipar et al., 2010) and natural infection (Herrewegh et al., 1997) indicates that the gastrointestinal (GI) tract may be a major site for FCoV persistence. Therefore, the study of viral diversity during persistent infections is essential to understand how viral evolution over time can result in viral variants having a range of disease profiles, including the acute presentation of FIP. While not typically resulting in FIP-like disease studies of sub-acute presentations of FCoV-1 may be informative in comparing natural infections of SARS-CoV-2, especially in immunocompromised humans (Corey et al., 2021); see (Pirofski & Casadevall, 2020) and Discussion.

In the present study, we studied FCoV-1 intra-host viral diversity, to assess if persistent FCoV-1 infection in a naturally infected cat promotes viral genetic variation over time, focusing on a region of the genome closely related to pathogenesis (the S1/S2 region). We used two approaches. First, we followed the cat for six years and collected mostly non-invasive samples (feces, conjunctiva, and nasal swabs, blood) to detect and characterize the diversity of FCoV-1 variants shed by the cat. Second, after the euthanasia of the cat, we collected 18 tissues and biofluids to study the viral diversity and distribution in the body of a persistently infected cat. Our results show that a cat without typical signs associated with FIP can have modified S1/S2 cleavage sites in various tissues and secretions—indicating a persistent GI reservoir for the virus and allowing the modulation of spike protein function leading to altered pathogenicity and transmission properties, including in the heart and conjunctiva.

## Methods

### Sample collection

A 5-year-old neutered male Bengal cat was enrolled in 2017 as a healthy cat in our long-term FCoV project (cat ID #576). Over this time period, the cat was clinically diagnosed with inflammatory bowel disease (IBD) and chronic rhinitis, with biopsies and histopathology performed in 2017. He was also diagnosed with a low-grade heart murmur secondary to mitral valve regurgitation. From 2017 to 2022, we collected various non-invasive samples, including feces and conjunctival, oro-pharyngeal, and saliva swabs (Table 1). In 2017, samples such as blood, and nasal and intestinal biopsies were taken during routine vet visits. The fecal, swab and blood samples were analyzed immediately after collection, and then frozen at -80°C until further use. The tissues resulting from the biopsies were preserved in formalin-fixed paraffin-embedded (FFPE) blocks by the Animal Health Diagnostic Center (AHDC), Cornell University. The samples taken in 2019 and 2020 were stored in DNA/RNAshield (Zymo Research) and submitted to the Metacats project (http://metasub.org/metacats). In 2022, the cat was euthanized at 11 years of age due to respiratory distress. Necropsy was performed by the Cornell University College of Veterinary Medicine pathology service. Eighteen tissues and biofluids were collected at necropsy and were deposited in the Cornell Veterinary Biobank (https://www.vet.cornell.edu/departments/centers/cornell-veterinary-biobank) under accession number 29801. The tissues were flash-frozen in liquid nitrogen and stored at -80°C until further use, the biofluids were frozen at -80°C immediately after collection.

**Table 1.** Samples from cat #576 collected and analyzed in this study.

| **Sample type** | **Collection date** | **FCoV-1 screening result** |
| --- | --- | --- |
| Nasal Biopsy | 04/28/17 | **Positive** |
| Intestinal Biopsy | 04/28/17 | **Positive** |
| White blood cells | 11/17/17 | Negative |
| Conjunctival swab | 11/17/17 | **Positive** |
| Oro-pharyngeal swab | 11/20/17 | Negative |
| Feces | 11/20/17 | **Positive** |
| Feces | 09/05/19 | Negative |
| Feces | 2020 | Negative |
| Saliva swab | 2020 | Negative |
| Brain | 01/06/22 | Negative |
| Lung | 01/06/22 | Negative |
| Urine | 01/06/22 | Negative |
| Feces | 01/06/22 | **Positive** |
| Stomach content | 01/06/22 | Negative |
| Kidney | 01/06/22 | Negative |
| Intestine - Ileum | 01/06/22 | Negative |
| Intestine - duodenum | 01/06/22 | Negative |
| Intestine - jejunum | 01/06/22 | **Positive** |
| Spleen | 01/06/22 | Negative |
| Heart ventricle | 01/06/22 | **Positive** |
| Liver | 01/06/22 | Negative |
| Pancreas | 01/06/22 | Negative |
| Lymph node | 01/06/22 | Negative |
| Eye, cornea | 01/06/22 | Negative |
| Mucosal tissue, nasal cavity | 01/06/22 | Negative |
| Plasma, EDTA | 01/06/22 | Negative |
| Serum | 01/06/22 | Negative |

### Viral screening and genetic analyses

Total RNA was obtained from tissues, feces and swabs using the Monarch Total RNA Miniprep Kit (New England Biolabs) adding a DNase step to deplete host DNA contamination and following manufacturer’s instructions for each type of sample. Before the RNA extraction, between 20 to 40 mg of sample was disrupted using the Bead Ruptor Elite (OMNI International) using the MagMAX Microbiome Bead Tubes for the fecal samples and 500mg of ∼2.5mm Zirconia/Silica beads (Biospec Products) for tissues and 650ul of Protection Reagent for all sample types. FCoV-1 RNA was detected by hybridization capture with a Twist Custom Panel (Twist Biosciences) targeting the S gene of UU4 and Black (accession numbers FJ938054 and EU186072, respectively). cDNA was synthetized from the extracted RNA using Superscript III Reverse Transcriptase (Thermo Fisher Scientific), followed by dsDNA synthesis using DNA Polymerase I, Large (Klenow) Fragment (New England Biolabs, NEB). Resulting dsDNA was cleaned using the Agencourt AMPure XP magnetic beads (Beckman Coulter). dsDNA was quantified using the Qubit dsDNA Broad Range Assay kit (Thermo Fisher Scientific). Libraries were prepared from the resulting dsDNA using the Twist Library Preparation EF Kit 2.0 (Twist Biosciences) and the Twist CD Index Adapter Set 1-96 (Twist Biosciences), according to manufacturer’s protocol. The hybridization capture was carried out using the Twist Hybridization and Wash Kit, the Twist Wash Buffers, and the Twist Universal Blockers (Twist Biosciences), following manufacturer’s protocol. The capture libraries were sequenced in the iSeq 100 using the i1 Flow Cell and Reagent Cartridge v2 (Illumina) for 300 cycles. Obtained reads were analyzed in Geneious Prime ® 2022.1.1. Reads with low quality (QC <20) were trimmed using the BBDuk algorithm. Trimmed reads were mapped against the sequence of the S gene of FCoV-1 UU4 and Black using the Geneious mapper using the medium sensitivity methods.

The obtained nucleotide (nt) sequences for all the positive samples were aligned using the Clustal Omega algorithm (Sievers et al., 2011). A haplotype network of the S1/S2 region (17 amino acids, 51nt) for seven variants was constructed using the median-joining algorithm in Network 5.0 (Sievers et al., 2011). We also aligned our nucleotide and amino acid sequences with 39 others available at Genebank of which the biotype (FECV, FIPV) was known, as previously published (Zehr et al., 2023).

### RNA-based *in situ* hybridization (ISH)

A probe targeting the RdRp gene of FCoV was used to detect viral RNA (RNAscope ® 2.5 VS Probe V-FIPV-ORF1a1b, Advanced Cell Diagnostics) using ISH (Sweet et al., 2022). The ISH process was carried by the AHDC using the automated staining platform Ventana Discovery Ultra (Roche Tissue Diagnostics) using the Discovery kits for mRNA Sample Prep, mRNA Red Probe Amplification and mRNA Red Detection and the Ventana Hematoxylin and Ventana Bluing Reagents for counterstaining.

### Tertiary structure prediction

The amino acid sequences of two variants obtained in 2017 (FCoV-1 feces 2017 and FCoV-1 conjunctiva 2017), were aligned with FCoV-1 UU4 spike protein, from which the tertiary structure has been solved by cryo-EM (PDB ID 6JX7) (Yang et al., 2020), using the Clustal Omega algorithm in Geneious Prime ® 2022.1.1. Using this alignment, the complete or partial S tertiary structure from our two variants were modelled in Swiss-Model (www.swissmodel.expasy.org) using the target-template alignment input. The two models were superimposed, and colors were modified using UCSF ChimeraX (Pettersen et al., 2021).

## Results

In total, we screened 27 samples (6 in 2017, 1 in 2019, 2 in 2020, and 18 in 2022) for the presence of FCoV-1 RNA, of which 7 were positive (Table 1). From the samples screened in 2017, we found FCoV-1 RNA in feces and the conjunctival swab, and in nasal and intestinal biopsy, and from the 18 samples screened in 2022 we found it in feces, heart, and intestine (jejunum). We did not find FCoV-1 RNA in the samples taken in 2019 (1 sample) or 2020 (2 samples).

After removing the low-quality reads from the seven positive samples, we were able to obtain the whole S1/S2 region from all samples (114 to 120nt, 37 to 40 amino acids), which includes the complete FCS (40 to 51nt, 15 to 17 amino acids), except from that from the intestine (jejunum) taken in 2022 from which we were only able to obtain a partial sequence that only includes a minimal FCS (6 amino acids, Figure 1A). Genetic analysis of the reads obtained from the heart in 2022 revealed that there were two single nucleotide variants (named heart1 and heart2) that differed in one nucleotide (T in heart1, A in heart2) in codon 14 of our alignment, which resulted in a non-synonymous change (arginine (R) in heart1, serine (S) in heart2) in the FCS (Figure 1A). No other single nucleotide variant was detected in the other sequences obtained. Therefore, we obtained the complete S1/S2 region of seven variants (feces, conjunctiva swab, nasal and intestine biopsies from 2017, and feces, heart1, and heart2 from 2022) and one partial sequence (intestine 2022). Nucleotide comparison of the whole S1/S2 region, from our 5 variants showed that there was size variation (120nt in conjunctival swab 2017, and 114nt in the other six variants, Figure 1A). This size difference is due to the insertion of 6 nucleotides in the downstream P’ location of the FCS, that results in two additional amino acids in the sample from the conjunctival swab 2017 when compared to the other six variants (Figure 1A). A search in the Genebank revealed that 10 additional FCoV-1 variants have insertions of different sizes in that same region resulting in 1 to 4 additional amino acids when compared to ours (Figure 2). Six of these were from FIPV variants and 4 from FECV variants.

**Figure 1.**
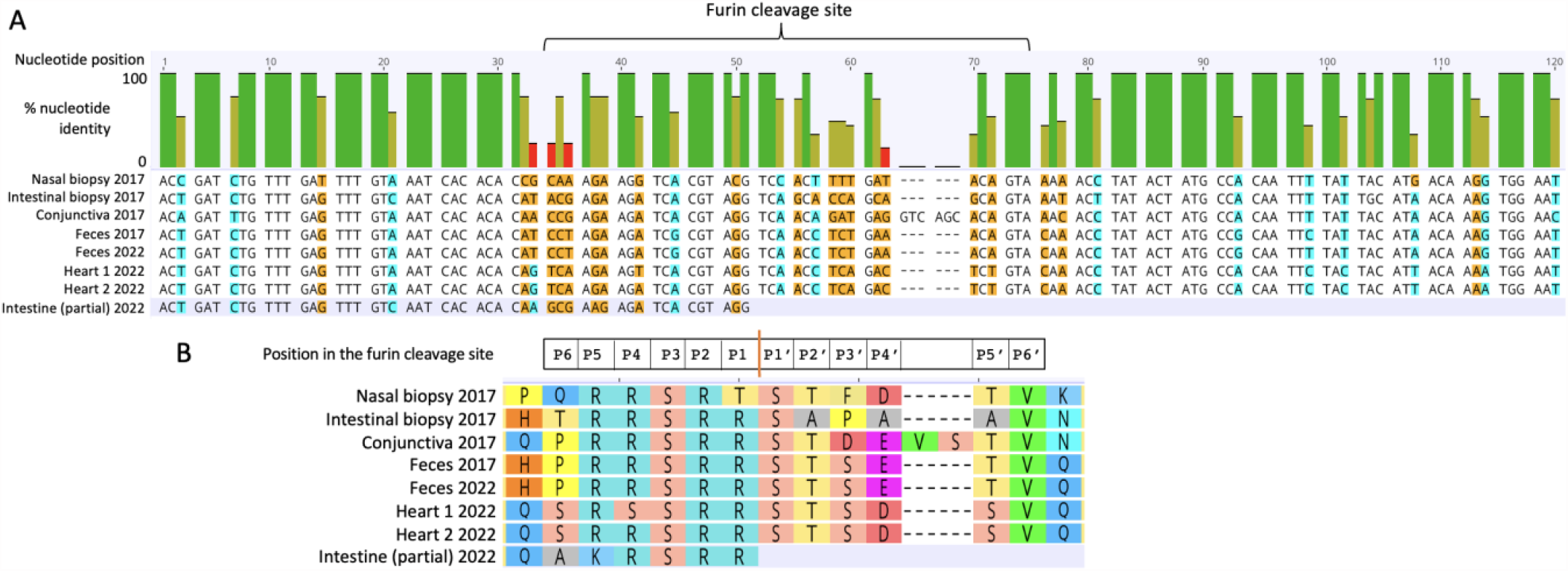
The S1/S2 region of the eight variants obtained in this study in 2017 and 2022. A. Codon alignment of the S1/S2 region (120nt long), including the furin cleavage site (FCS). The top graph shows the percentage of nucleotide sequence identity for each site. In the nucleotide alignment, nucleotides highlighted in cyan indicate synonymous mutations (*i.e*. those that do not result in a change in amino acid), while those in orange are non-synonymous mutations (*i.e*. those that result in an amino acid change. A hyphen indicates a deletion. B. The corresponding amino acids for each codon of the FCS shown in A. Above the amino acid alignment are the P6 to P6’ positions of each amino acid in the FCS. The site in which the cleavage occurs is shown as a vertical orange line.

**Figure 2.**
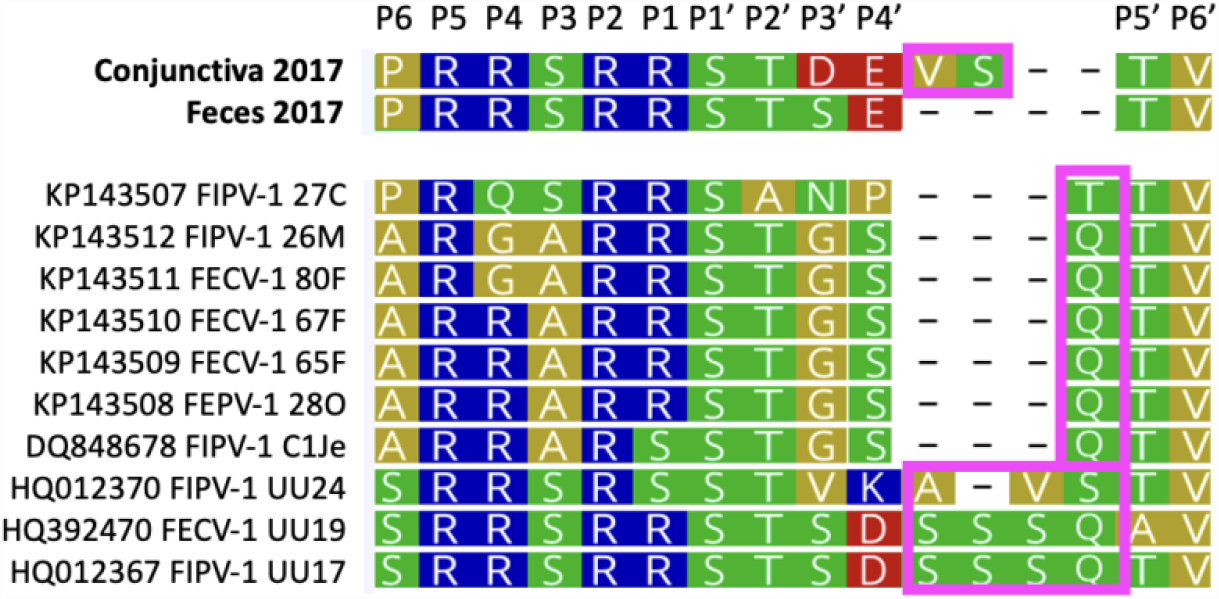
Amino acid alignment of the S1/S2 cleavage site of two FCoV-1 variants obtained in this study (on top, in bold) and 10 variants available on NCBI showing various amino acid insertions in the downstream P’ location (enclosed in a pink square). The position of the residues involved in the FCS (P6 to P6’) is at the top of the alignment and is based on the shortest reference variants obtained in this study (feces 2017). The ten sequences obtained from NCBI have one to four amino acid insertions in the same downstream P’ location as the conjunctiva 2017 variant obtained in this study (enclosed in a pink square). The accession numbers for all these ten sequences are shown.

A comparison of the nucleotide sequences of the S1/S2 region revealed that there was intra-host nucleotide variation in this region. In the 120nt region sequenced there were mutations in 34 positions (in orange and cyan in Figure 1A). Most mutations that result in a change of residue (non-synonymous, in orange in Figure 1A) were within the FCS (19 out of 21 non-synonymous mutations) resulting in a variety of different residues (Figure 1B). From the seven complete FCS amino acid sequences obtained, only those from feces in 2017 and 2022 were identical. The four sequences obtained in 2017 differed in positions P6, P1, P2’, P3’, P4’ and P5’ and the four obtained in 2022 (including the partial intestine 2022) differed in P6, P5, P4, P4’, and P5’ (Figure 1B). Tertiary structure prediction of the complete S protein of variants obtained from feces in 2017 and the partial S1/S2 sequence of conjunctiva from 2017 (Figure 3A) showed that the S1/S2 loop of the variant obtained from conjunctiva in 2017 protrudes more than that of the feces 2017 (Figure 3B), possibly due to the two extra amino acids that it has (Figure 1B).

**Figure 3.**
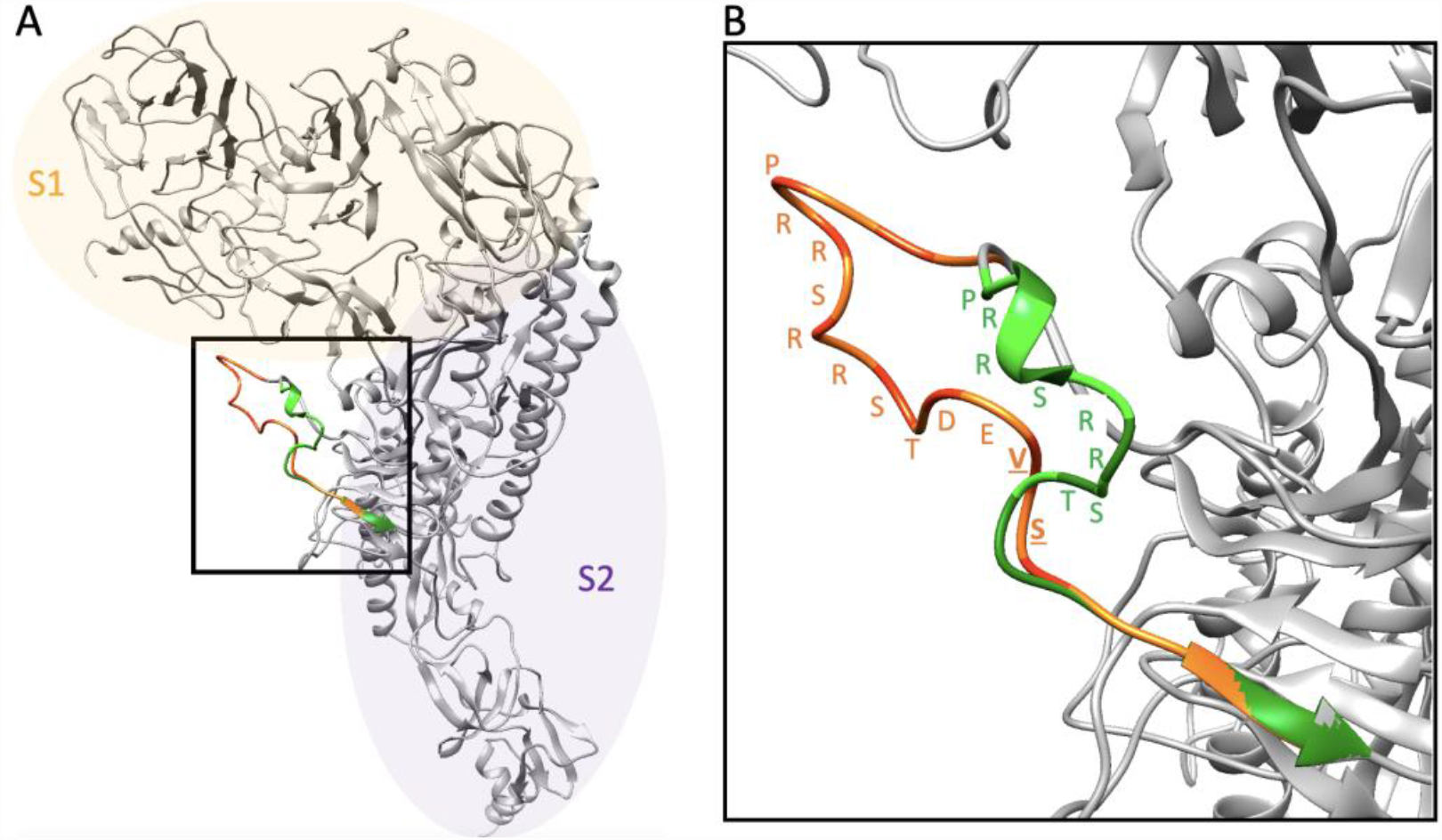
A. Tertiary structure prediction of the complete S protein of the FCoV-1 recovered from feces in 2017 (in gray). **The S1/S2 loop (inside the square) of this variant is highlighted in green and** superimposed to it is the partial S sequence obtained from conjunctiva in 2017 (in orange). **B. Closeup of the superimposed S1/S2 loop of the FCoV-1 variants from feces (in green) and conjunctiva (in orange) recovered in 2017 showing the amino acid sequence of the S1/S2 cleavage motif**. Underlined are the two additional residues observed in the variant obtained from conjunctiva.

A haplotype network based on the complete nucleotide sequence of the FCS from seven variants revealed that the sequences did not group by year of collection but by site within the host (Figure 4). Both variants obtained from feces were identical and the variants from heart only differed in the one nucleotide reported above. The other three variants (nasal biopsy, intestinal biopsy, and conjunctiva 2017) were placed in different nodes with 6 to 9 nucleotide mutations to internal nodes. The internal nodes are connected in three intermediate loops. Mapping of the nucleotides in each branch of the loops shows that each contains mutations in codons from amino acids within the FCS (P6, P3’, P4’ in Figure 3).

**Figure 4.**
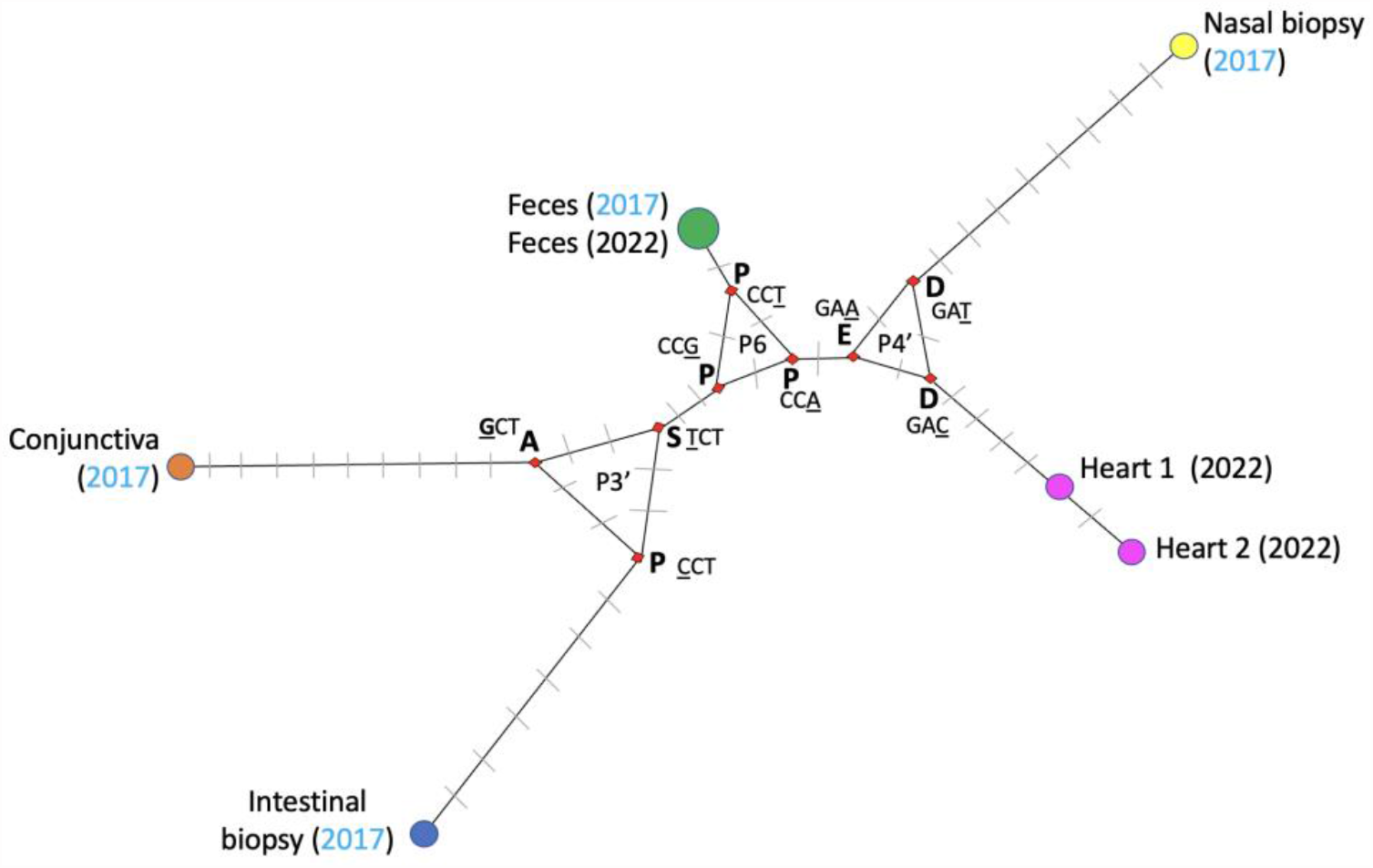
Haplotype network of the S1/S2 region (120nt) of seven variants of FCoV obtained from cat #576 in 2017 and 2022. Circles represent haplotypes (or unique sequences); the size of each circle is proportional to the number of variants within each node. Next to each circle is the collection place and year (in parenthesis) of the given haplotype. The color of the circles indicates the sample type: blue is intestine, orange is conjunctiva, pink is heart, green is feces, yellow is nasal cavity. The length of each branch is proportional to the number of nucleotide changes between haplotypes and gray lines on branches indicate the nucleotide changes between nodes or haplotypes. Red squares indicate intermediate nodes. Residues mapped for each intermediate node are shown next to the node in bold in single-letter code. Next to each residue is the corresponding codon and the nucleotide mutation in each branch is underlined. Inside each intermediate loops are the positions within the FCS in which the mutations are occurring and include P3’, P6, and P4’.

To examine the possible tissue distribution of FCoV-1, samples from necropsy were examined by histopathology, using a newly developed RNA-in-situ hybridization technique (Sweet et al., 2022), with the goal of improving sensitivity in the possible sites of viral persistence. Using this technique, FCoV RNA was detected in isolated cells in the heart, and in epithelial cells of the small intestine (jejunum) (Figure 5).

**Figure 5.**
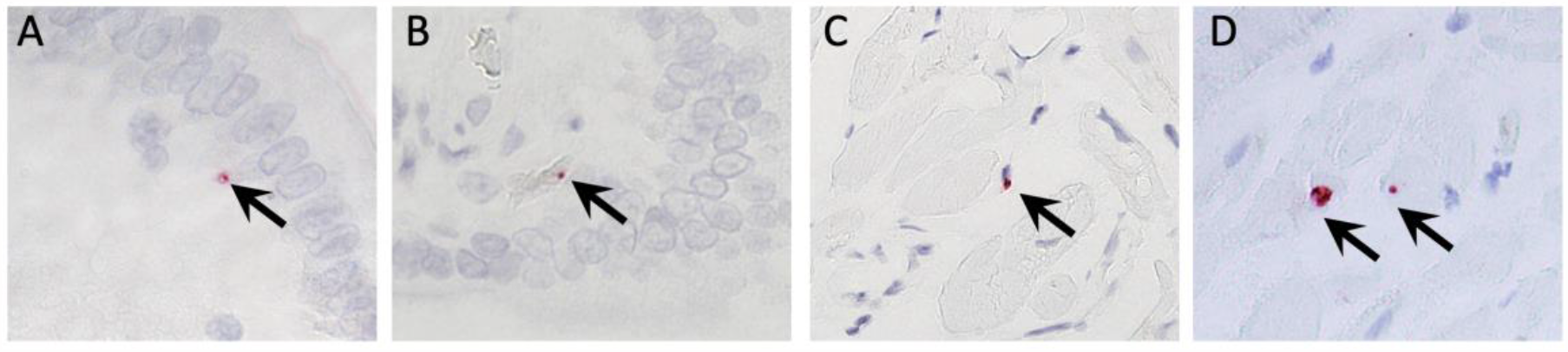
Detection of FCoV RNA (in red, indicated by arrows) by in-situ hybridization in (A, B) intestine and (C, D) heart of cat #576 in 2022. All photos were taken at x600 magnification.

## Discussion

In this study, we provide information on persistent coronavirus infection in a naturally infected cat with FCoV-1. We show that FCoV-1 variants detected in the feces of the same individual 5 years apart (in 2017 and 2022) had an identical FCS sequence (Figure 1). Our results together with previous reports of long-term shedding of highly conserved FCoV-1 (>97% nucleotide similarity) in feces (Addie et al., 2003) support the idea that continuous viral shedding by an individual is the result of FCoV-1 persistent infection, rather than being a re-infection with different variants. By taking multiple samples from an infected individual during a 5-year period and using next-generation sequencing we were also able to characterize the anatomical distribution of FCoV-1 during a persistent natural infection. In line with previous studies (Herrewegh et al., 1997) we show that FCoV-1 can persist in the intestine, but we also show that it can persist in sites outside the gastrointestinal system like the heart (Figure 1). Additionally, during persistent infection, FCoV-1 can also be detected in the nasal cavity and conjunctiva (Figure 1). Viral persistence is often associated with viruses residing in immune-privileged sites like the eyes and testes, in which for example, viruses like Ebola and Zika have been reported (Dyal et al., 2022; Pierson & Diamond, 2018). In this study, we show that FCoV-1 RNA persistence is not restricted to sites classically considered immune privileged (*i.e*., we did not detect FCoV-1 RNA in the brain or the eyes, Table 1), but it can persist in the heart and intestine (Figure 1).

We also show that a cat persistently infected with FCoV-1 harbors different viral subpopulations in different tissues. Although we found that the variants shed in the feces were identical, they were different from those detected in tissues (heart, intestine, nasal cavity) and fluids (conjunctiva, Figure 4). Additionally, different tissues harbored variants of FCoV-1 that differed in the sequence of the S1/S2 FCS (Figures 1 and 4). FCoV-1 RNA can be detected in various organs after experimental infection (Kipar et al., 2010) even if the cats do not develop FIP (Lutz et al., 2020). Likewise, FCoV-1 has been detected in different tissues and feces in 24% of cats that died of unrelated illnesses (Jähne et al., 2022) Experimental infection also suggests that shortly after infection FCoV-1 undergoes continuous but subtle changes, especially in the S gene (Desmarets et al., 2016) and that the composition of the viral quasispecies varies among different organs, although the sequences were not obtained (Kiss et al., 2000). This is similar to what has been reported for SARS-CoV-2, for which tissue-specific high-frequency variants outside the pulmonary system have been detected within the first 10 days after the first detection of the virus in humans (Normandin et al., 2023). Considering this, our hypothesis is that after initial infection, FCoV-1 spreads to different tissues where it replicates at varying rates and results in the emergence of different variants in each tissue. Some of these variants may persist in certain tissues and will remain genetically stable over time (*i.e*., the variants from feces taken five years apart were identical).

Interestingly, associated pathologies related to the tissues in which we found FCoV-1 (intestine, nasal cavity, heart) were reported for this cat and included a long history of IBD and heart murmur. The cat also suffered from chronic rhinitis, and although we found FCoV-1 RNA in the nasal biopsy taken in 2017, we were not able to find FCoV-1 RNA in the mucosal tissue of the nasal cavity taken in 2022 (Table 1). Therefore, we are not able to determine whether the virus remained in the nasal cavity over time. In humans, persistent infection with different viruses has also been related to chronic symptoms that linger after the patient’s recovery from the acute infection. For example, cardiomyopathies have been related to persistent infection with enterovirus (Kühl et al., 2005) and SARS-CoV-2 (Omidi et al., 2021). Chronic clinical symptoms like weakness, fatigue, memory loss, and ataxia have also been linked to persistent viral infection with West Nile Virus (WNV) (Murray et al., 2010), and SARS-CoV-2 where it is also known post-acute sequelae of coronavirus infection, PASC, or “long covid” (Halpin et al., 2021). Although further studies are needed to assess if by establishing a viral reservoir in different tissues, FCoV-1 can cause tissue-specific pathologies (*i.e*., IBD, heart murmur, rhinitis), several studies have linked FCoV infection with certain chronic ailments of cats. For example, myocarditis has been previously reported in an FIP cat (Ernandes et al., 2019) with pronounced viral antigen in the heart detected by IHC, along with a report of FIP-linked dilated cardiomyopathy (Yoshida et al., 2016). There are also reports of FCoV in the heart tissue/muscle in a sub-clinical cat (Kiss et al., 2000) and one with epicarditis (Araujo et al., 2020). In addition, (Malbon et al., 2019) noted the involvement of both the liver and heart in FCoV pathogenesis. However, no direct assessment of FCoV infection of heart tissue was evaluated, either by PCR or histology. FCoV-1 infection has also been detected in a cat with rhinitis (Andre et al., 2020) and in persistently infected cats with chronic diarrhea (Addie & Jarrett, 2001). It is also important to note that a recent report revealed that administering the non-FDA-approved anti-viral drug GS-441524 to cats infected with FCoV-1 and experiencing chronic enteropathy or IBD resulted in the elimination of viral shedding in feces and the resolution of chronic gastrointestinal symptoms in all cats involved in the study (n=7) (Addie et al., 2023).

Here we also provide evidence that a persistently infected cat can shed FCoV-1 RNA intermittently in the feces for at least five years. These results show that even if a cat does not display typical signs of FCoV-1 infection, by shedding the virus in feces and conjunctiva, this individual is epidemiologically relevant as it may spread the virus to other cats. This is of particular interest as previous studies have shown that the majority (up to 80%) of naturally and experimentally infected cats remained persistently infected without showing signs typically associated with FCoV-1 infection and constantly shed the virus in feces (Kiss et al., 2000). Therefore, cats that are persistently infected and shed the virus (intermittently or constantly) may play a crucial role in maintaining the high prevalence of FCoV-1 in multi-cat environments, including closed cat-breeding facilities (Herrewegh et al., 1997). We also found FCoV-1 in the intestine in 2017 and 2022, however, we were only able to get a partial sequence from the variant taken from the intestine (jejunum) in 2022 (Figure 1). In healthy cats infected with FCoV-1, it has been reported that the virus is restricted to a single or a few cells in the mucosal surface of the colon (Kipar et al., 2010). It has also been reported that during the course of the infection, FCoV-1 is detected in different sections of the intestine (colon, ileum, duodenum, jejunum) in a single individual. This suggests that viral load in the intestine fluctuates over time, which could explain why we were able to sequence the complete FCS of the variant from the intestine in 2017 but not from 2022 (Figure 1). This is also supported by the detection by ISH of FCoV RNA in very few cells in the intestine sample taken in 2022 (Figure 5), indicating that the viral load was minimal. As this (Figure 5) and a previous study (Kipar et al., 2010) detected FCoV-1 in superficial epithelial cells of the intestine, an intestinal biopsy that mainly removes superficial cells could be a viable sample option for detecting FCoV-1. This could be helpful for cats suffering from chronic GI diseases that may need an intestinal biopsy for a complete diagnosis.

In addition, we demonstrate that there is variation in the FCS of FCoV-1 not only through sequence mutations but also through insertions in the downstream P’ region of the FCS (Figure 1). Based on the structural model presented in Figure 3, we suggest that the two extra amino acids observed in the variant from the conjunctiva 2017, result in a long and more exposed loop which may be more accessible to the furin protease. Equivalent insertions consisting of 1 to 4 additional amino acids have also been documented in the same location in other FCoV-1 spikes (Figure 2). While these insertions do not appear to correlate specifically with FECV or FIPV they may correlate with a history of enhanced shedding, and may have been retained when the virus advanced to FIP through other genomic mutations, including in S2’ (Licitra et al., 2014). For example, the first six variants in Figure 2, which have an additional amino acid, were similarly detected during an ongoing outbreak of FIP (Lewis et al., 2015). Equivalent insertions or deletions in the downstream region of the FCS have recently been documented in other coronavirus spike proteins. Notably, an HCoV-OC43 with a 4 amino acid insertion was reported in a patient with fatal pneumonia (Lau et al., 2021). HCoV-OC43 typically infects the upper respiratory tract. However, this variant was found in high viral loads in the lower respiratory tract (Lau et al., 2021). It remains to be determined if this amino acid insertion is linked to the presence of a novel circulating genotype or with the observed change in tissue tropism. While such P’ indels have not, to our knowledge, been seen with SARS-CoV-2 spike, the Delta variant with increased furin cleavability (in this case linked primarily to its P681R mutation in the FCS) fits this model; see refs (Liu et al., 2022; Lubinski et al., 2022; Saito et al., 2021) for examples. Our hypothesis is that for FCoV-1, adding amino acids near the FCS may increase FCoV-1 spike S1/S2 cleavage by further exposing the S1/S2 loop. This potentially enhances virus transmissibility through the respiratory tract or conjunctival routes.

To the contrary—and in the case of FIP—tissue/macrophage-specific variants of FCoV-1 are widely considered to be less transmissible. This situation may also be paralleled for of SARS-CoV-2, where loss of transmissibility seems to occur readily; *e.g*. with Vero cell-adapted isolates via both deletions and point mutations in the FCS region itself (Klimstra et al., 2020; Lamers et al., 2021; Sasaki et al., 2021), and with the upstream QTNTN motif that when deleted results in a shortened, more rigid peptide loop and less accessibility to protease (Vu et al., 2022). This study also shows that several sequence changes occur within the FCS in various tissues and fluids from a single host in a cat without a clinical diagnosis of FIP (Figure 4). In the heart, we found two variants that differed in their FCS sequences. Variant heart2 had the canonical furin cleavage motif (R-R-S-R-R|S) while variant heart1 had a disrupted motif due to an S in position P4 (R-S-S-R-R|S, Figure 1). Furin cleavage assays have shown that an R to S mutation in P4 in the FCS of FCoV-1 results in full abrogation of furin cleavage (Licitra et al., 2013). Therefore, our prediction for the heart1 variant (Figure 1), is that since the FCS is disrupted, the virus has switched the entry pathway to a cathepsin-primed route (as is expected with Vero cell-adapted SARS-CoV-2). We consider that the heart1 variant has lost its transmissibility potential, with the additional outcome that virus infection would result in reduced cell-cell fusion and less overt clinical signs in heart tissue/cardiomyocytes. In the case of SARS-CoV-2, loss of transmissibility seems to occur readily; *e.g*. with Vero cell-adapted isolates via both deletions and point mutations in the FCS region itself (Klimstra et al., 2020; Lamers et al., 2021; Sasaki et al., 2021). Our study suggests that FCS down-regulation may lead to a “silent” or quiescent infection of the heart tissue, in which the virus can persist over time. Therefore, FCoV-1 FCS genetic variation in certain tissues could be the result of viral adaptation to cause less damage to be able to persist over time in a host, as has been reported for SARS-CoV-2 in humans (Badrinath et al., 2022; Clemens et al., 2023; Navaratnarajah et al., 2021). Overall, our study of cat #576 leads to a new proposed model of FCoV-1 infection, depicted in Figure 6.

**Figure 6.**
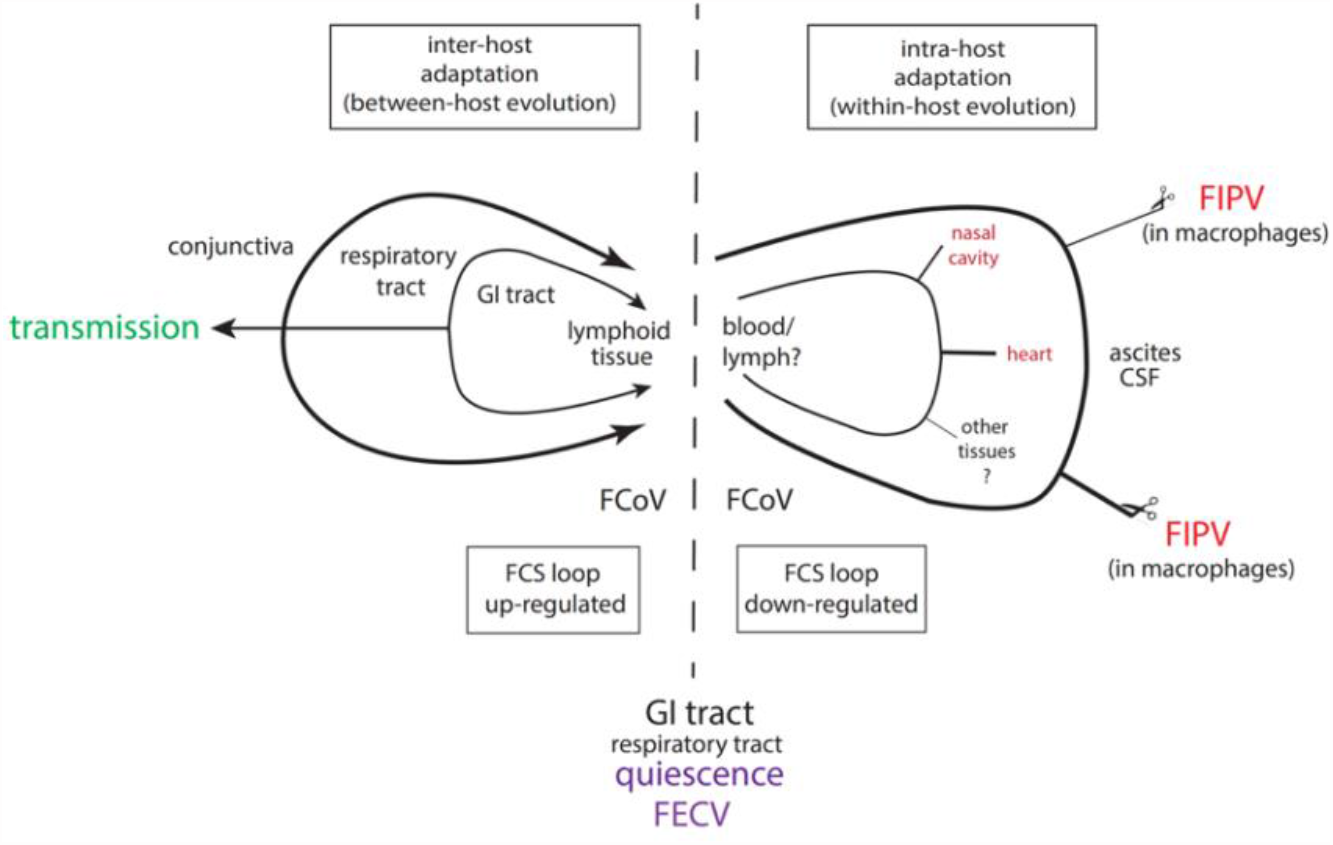
Proposed model of FCoV-1 dynamics, including data from cat#576.

## Acknowledgments

This work was supported by the Cornell Feline Health Center and by the Michael J. Zemsky Fund for Feline Disease at Cornell University College of Veterinary Medicine. Research reported in this publication was also supported by the National Center of Advancing Translational Science of the National Institutes of Health under Award Number UL1TR002384.

